# ALDH9A1 Deficiency as a Source of Endogenous DNA Damage that Requires Repair by the Fanconi Anemia Pathway

**DOI:** 10.1101/2022.05.26.493623

**Authors:** Moonjung Jung, Isaac Ilyashov, Yeji Park, Frank X. Donovan, Ramanagouda Ramanagoudr-Bhojappa, Danielle Keahi, Jordan A. Durmaz, Haruna B. Choijilsuren, Audrey Goldfarb, Mia Stein, Jungwoo Kim, Ryan R. White, Sunandini Sridhar, Raymond Noonan, Tom Wiley, Thomas S. Carroll, Francis P. Lach, Arleen D. Auerbach, Ileana Miranda, Settara C. Chandrasekharappa, Agata Smogorzewska

## Abstract

Fanconi anemia (FA) pathway is required for the repair of DNA interstrand crosslinks (ICL). ICLs are caused by genotoxins, such as chemotherapeutic agents or reactive aldehydes. Inappropriately repaired ICLs contribute to hematopoietic stem cell (HSC) failure and tumorigenesis. While endogenous acetaldehyde and formaldehyde are known to induce HSC failure and leukemia in humans with FA, the effects of other toxic metabolites in FA pathogenesis have not been systematically investigated. Using a metabolism-focused CRISPR screen, we found that ALDH9A1 deficiency causes synthetic lethality in FA pathway-deficient cells. Combined deficiency of ALDH9A1 and FANCD2 causes genomic instability, apoptosis, and decreased hematopoietic colony formation. *Fanca^−/−^Aldh9a1^−/−^* mice exhibited an increased incidence of ovarian tumors. A suppressor CRISPR screen revealed that the loss of ATP13A3, a polyamine transporter, resulted in improved survival of *FANCD2^−/−^ALDH9A1^−/−^* cells. These findings implicate high intracellular polyamines and the resulting 3-aminopropanal or acrolein in the pathogenesis of FA. In addition, we find that ALDH9A1 variants may be modifying disease onset in FA patients.

**Statement of Significance:** ALDH9A1 deficiency is a previously unrecognized source of endogenous DNA damage. If not repaired by the Fanconi anemia pathway, such damage leads to increased genomic instability and tumorigenesis. Limiting substrates that accumulate when ALDH9A1 is absent can reduce aldehyde production and rescue synthetic lethality in the combined deficiency of ALDH9A1/FANCD2.

## Introduction

Fanconi anemia (FA) is the most common inherited bone marrow failure (BMF) syndrome^1,2^. Up to 90% of patients develop BMF by the age of 40, with a high frequency of disease progression to myelodysplastic syndromes and acute myeloid leukemia^1,3^. Currently, allogeneic hematopoietic stem cell transplantation is the only curative treatment for HSC failure in FA. However, it still poses significant risks of early mortality and it increases the already high risk of solid tumors, including squamous cell carcinomas of the aerodigestive tract^4,5^. FA is caused by mutations in one of the 22 known *FANC* genes (*FANCA – FANCW*)^6–8^. Cells from patients with FA exhibit increased chromosomal breakage when exposed to crosslinking agents, such as mitomycin C (MMC) or diepoxybutane (DEB), which create DNA interstrand crosslinks (ICL)^9^. When ICLs are not properly repaired due to the failure of the FA pathway, HSCs activate the p21-p53 pathway and undergo cell cycle arrest and apoptosis^10^.

While the cellular defect in FA is impaired DNA ICL repair, the source of the DNA damage itself is not well understood. Recent studies in genetically modified mouse models combining deficiency of the aldehyde detoxification system and the FA pathway deficiency point to acetaldehyde and formaldehyde as important endogenous aldehydes that create ICLs^11–13^. Detoxifying enzymes – acetaldehyde dehydrogenase 2 (ALDH2) and alcohol dehydrogenase 5 (ADH5) – are required to metabolize these reactive aldehydes and prevent stem cell failure and leukemia in FA^14^. These findings are corroborated by human studies showing that dominant-negative *ALDH2*2* variants in FA patients^15^ and combined *ADH5/ALDH2* deficiency both result in accelerated bone marrow failure and leukemia ^16–18^.

Humans are exposed to more than 300 aldehyde compounds produced endogenously or found in the environment, including food^19^. There are 19 *ALDH* and 8 *ADH* superfamily genes in humans, working redundantly to metabolize these aldehyde species^20^. Despite extensive research, the complex defense network of aldehyde detoxification is yet to be fully understood.

To discover previously unknown endogenous sources of ICLs we performed a CRISPR knockout screen comparing survival of the wild-type (WT) and *FANCD2^−/−^* Jurkat cells. We found that the loss of ALDH9A1 cause synthetic lethality in *FANCD2^−/−^*cells, due to DNA damage-induced apoptosis. While the degree of DNA damage caused by combined deficiency of ALDH9A1 and FA pathway was not enough to cause spontaneous BMF in a mouse model, it led to an increased incidence of solid tumors with aging. Loss of ATP13A3 rescued the synthetic lethality of *FANCD2^−/−^ALDH9A1^−/−^*cells most likely through lowering substrates for ALDH9A1 and decreasing DNA damage.

## Results

### Identification of ALDH9A1 as a protein necessary for the survival of FANCD2 deficient cells

We hypothesized that a knockout of a pathway that would increase a metabolite capable of creating ICLs, necessitating the Fanconi anemia pathway function, would result in death or slow growth of *FANCD2^−/−^* cells. To identify novel endogenous sources of DNA damage, we performed CRISPR knockout (KO) screen using a metabolism-focused sgRNA library (3,000 genes, 10 sgRNA per gene, total 30,000 sgRNA) in a Cas9-expressing lentiviral vector^21^ in both WT and *FANCD2*^−/−^ Jurkat cells (Figure 1A). *FANCD2*^−/−^ Jurkat clones used in our experiments were created by transient expression of Cas9-sgRNA lentiviral vector followed by single-cell cloning. FANCD2 protein expression was confirmed to be absent in four KO clones with biallelic *FANCD2* mutations, and two selected clones were used for experiments (Figure 1B-D; S1A-D). They were sensitive to mitomycin C, the hallmark of FA (Figure 1C). For the screen, WT and *FANCD2*^−/−^ Jurkat cells were infected at multiplicity of infection (MOI) of 0.3 in biological triplicates. 1000x representation per sgRNA was maintained throughout the experiment. sgRNA representation was assessed immediately after puromycin selection or after 14 population doubling.

**Figure 1.**
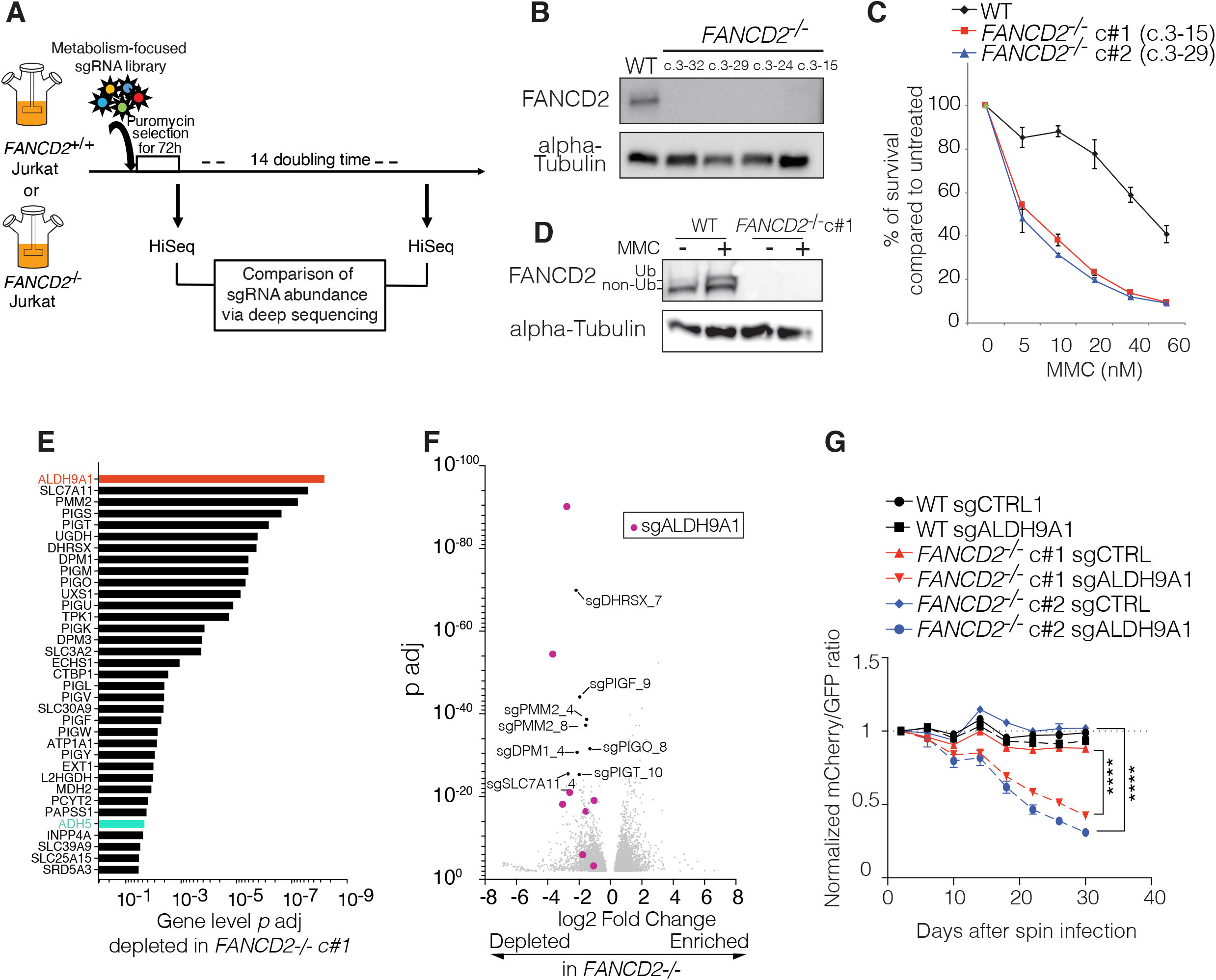
Metabolism-focused CRISPR screen reveals novel endogenous sources of DNA damage. A. A schematic summary of the metabolism-focused CRISPR knockout screen. WT and *FANCD2^−/−^* Jurkat cells were infected with lentivirus containing metabolism-focused sgRNA library in biological triplicates at MOI of 0.3. After 72 hours of puromycin selection, 40 million cells were harvested for baseline sequencing to achieve 1000x representation per sgRNA. At least 40 million cells were passaged every 4 days for 14 population doublings, after which at least 40 million cells were harvested for final time point sequencing. B. FANCD2 western blot of Jurkat clones used in the experiments. C. MMC sensitivity assay of Jurkat clones. The experiment was performed in triplicates. Three independent experiments were performed using WT and *FANCD2^−/−^* c#1. The third experiment included *FANCD2^−/−^* c#2, which is shown in this figure (mean +/- SD). D. FANCD2 western blot of Jurkat clones with and without MMC 1uM 24-hour treatment. E. Adjusted *p (p* adj) values for top 35 genes that were differentially depleted in the *FANCD2^−/−^* condition were calculated by the DESeq2 package and shown as a minus log 10 graph. F. A volcano plot was generated with log2 fold change (log2FC) value and *p* adj between *FANCD2^−/−^* and WT at the sgRNA level. Only sgRNAs with *p* adj <0.01 were used to generate this volcano plot. Eight significantly depleted sgALDH9A1s were marked as purple dots. G. A competition assay using WT and *FANCD2^−/−^* clones. The ratio of mCherry/GFP normalized to baseline was decreased in two *FANCD2^−/−^* clones with mCherry-sgALDH9A1, compared with *FANCD2^−/−^* clones with mCherry-sgCTRL (non-targeting). There were no differences between mCherry-sgALDH9A1 and mCherry-sgCTRL in WT. One-way ANOVA followed by Tukey’s multiple comparison test was performed for the last time point between *FANCD2^−/−^* c#1 sgCTRL and sgALDH9A1, and between *FANCD2^−/−^*c#2 sgCTRL and sgALDH9A1. (****, *p*<0.0001). MOI, multiplicity of infection; MMC, mitomycin C; SD, standard deviation.

Results of the screen are shown in Table S1-2. *ALDH9A1* was the top gene identified as a novel synthetically lethal gene with *FANCD2* deficiency (Figure 1E; Table S1). Eight out of ten sg*ALDH9A1*s were significantly depleted in *FANCD2*^−/−^ Jurkat cells (Figure 1F; Table S2). A*DH5,* but not *ALDH2* was also significantly depleted in *FANCD2*^−/−^ Jurkat cells, indicating that the sensitivity to different aldehydes may depend on the cell type.

To validate ALDH9A1 as having synthetic lethal interaction with FA deficiency, we performed an *in vitro* fluorescence-based competition assay (see Methods). We found an increased loss of cells expressing mCherry-sgALDH9A1 in two independent *FANCD2^−/−^* Jurkat clones, but not in WT cells (Figure 1G). *FANCD2^−/−^* cells targeted by sgALDH9A1 also showed slower growth compared with *FANCD2^−/−^* cells targeted by sgCTRL (Figure S1E). Targeting of *FANCD2^−/−^* cells by sgADH5, but not by sgALDH2 or sgCTRL, showed similar results. Differences were not observed in WT cells (Figure S1F-G).

*SLC7A11* and *SLC3A2*, the cystine/glutamate antiporters which form System Xc-(also known as xCT) were the second and sixteenth hits in our screen. They provide reduced glutathione for redox balance and formaldehyde metabolism ^22–25^. While there was an increased depletion of sgSLC7A11s in *FANCD2^−/−^* cells in the pooled screen, the loss of SLC7A11 was also detrimental to the growth of WT cells (Figure S1H) indicating the essential nature of their function in Jurkat cells.

Multiple genes in the glycosylphosphatidylinositol-anchored protein (GPI-AP) biosynthesis pathway^26^, were found to be significantly depleted in *FANCD2*^−/−^ compared with WT Jurkat cells (Figure 1E). Gene Ontology (GO) enrichment analysis confirmed this conclusion (Figure S1I). As the proper transfer of GPI-AP to nascent protein serves as endoplasmic reticulum (ER) exit signal^26^, it is possible that increased ER stress causes preferential cell death in *FANCD2^−/−^* cells.

### Combined deficiency of ALDH9A1 and FANCD2 results in increased DNA damage and apoptosis

To investigate the cause of synthetic lethality observed in ALDH9A1 and FANCD2 deficiency, WT and *FANCD2^−/−^* Jurkat cells were edited by ribonucleoprotein (RNP) delivery of Cas9 protein and sgALDH9A1 or sgCTRL by nucleofection (Figure 2A). With this approach, we achieved consistently high editing efficiency in both WT and the two independent *FANCD2^−/−^*clones at baseline. Sequencing on days 24 and 35 revealed a selective decrease in insertions/deletions (indel) levels only in the two *FANCD2^−/−^* clones, confirming a preferential loss of cells with combined FANCD2 and ALDH9A1deficiency, but not with ALDH9A1 deficiency alone (Figure 2B). Increased DNA damage was evident by the presence of increased chromosome breakage and increased numbers of 53BP1 foci and gamma-H2AX foci in *FANCD2^−/−^*, but not in WT cells targeted by sgALDH9A1 (Figure 2C-F, S1J). Cells with combined FANCD2 and ALDH9A1 deficiency also displayed increased Annexin-V+ cells and sub-G0 fraction on cell cycle analysis, consistent with DNA damage-induced apoptosis (Figure 2G-H).

**Figure 2.**
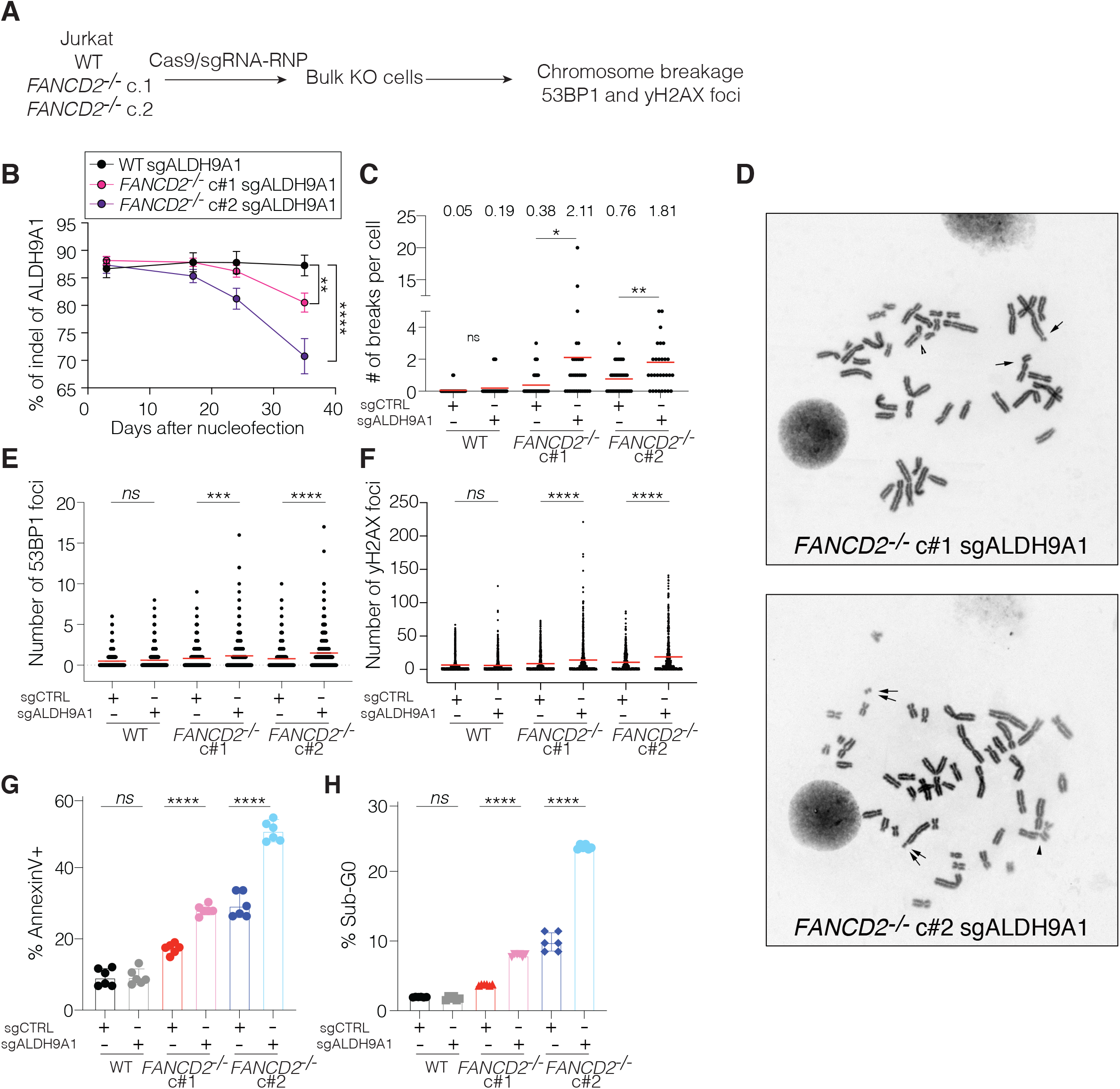
Loss of both ALDH9A1 and FANCD2 causes increased DNA damage due to endogenous insults and increased apoptosis. A. Schematic of bulk KO model. WT and two independent *FANCD2^−/−^*clones were targeted by sgALDH9A1 or sgCTRL (non-targeting) by electroporation of Cas9-sgRNA RNP complex. B. Percentage (%) of ALDH9A1 indel was assessed by Sanger sequencing followed by Synthego ICE analysis. ALDH9A1 indel (%) was decreased over time only in two independent *FANCD2^−/−^* clones, but not in WT. Four independent experiments were performed. One-way ANOVA followed by Dunnett’s multiple comparison test (**, *p*=0.0055; ****, *p*<0.0001). C. Numbers of chromosome breakage per cell were significantly elevated in two *FANCD2^−/−^* clones targeted by sgALDH9A1, compared with clones targeted by sgCTRL (non-targeting). There were no differences in WT clones between sgCTRL and sgALDH9A1. Student t test was performed between sgCTRL and sgALDH9A1 in each clone (*, *p*<0.05; **, *p*<0.005). D. Representative figure of chromosome breakage analysis. E and F. Numbers of 53BP1 foci per cell (E) and gamma-H2AX foci per cell (F) by immunofluorescence were elevated in two *FANCD2^−/−^* clones targeted by sgALDH9A1, compared with clones targeted by sgCTRL (non-targeting). Two independent experiments were performed. G and H. Percentage of Annexin V-positive cells (G) and sub-G0 fraction on cell-cycle analysis (H) by flow cytometry were elevated in two *FANCD2^−/−^*clones targeted by sgALDH9A1, compared with clones edited with sgCTRL (non-targeting). Two independent experiments were performed. E-H. One-way ANOVA followed by Tukey’s multiple comparison test was performed (*ns*, not significant; *, *p*<0.05; **, *p*<0.005; ***, *p*<0.0005;****, *p*<0.0001).

### Combined ALDH9A1 and the FA pathway deficiency decreases the survival of HSC

To examine the effect of combined deficiency of ALDH9A1 and the FA pathway on HSCs, we edited *FANCD2* and/or *ALDH9A1* in human umbilical cord blood CD34+ cells (Figure 3A). While cutting efficiency for each gene at baseline was lower when attempting a double KO (Figure 3B), the double KO cells produced the fewest hematopoietic colonies and the lowest frequency of GEMM colonies (Figure 3C-D). To examine the editing and zygosity of individual HSPC, each colony was harvested and sequenced. We observed fewer colonies with mutations both genes (biallelic double KO; observed to expected ratio 0.33) as compared to either single gene KO, suggesting decreased survival of HSC by combined deficiency of FANCD2 and ALDH9A1 (Figure 3E).

**Figure 3.**
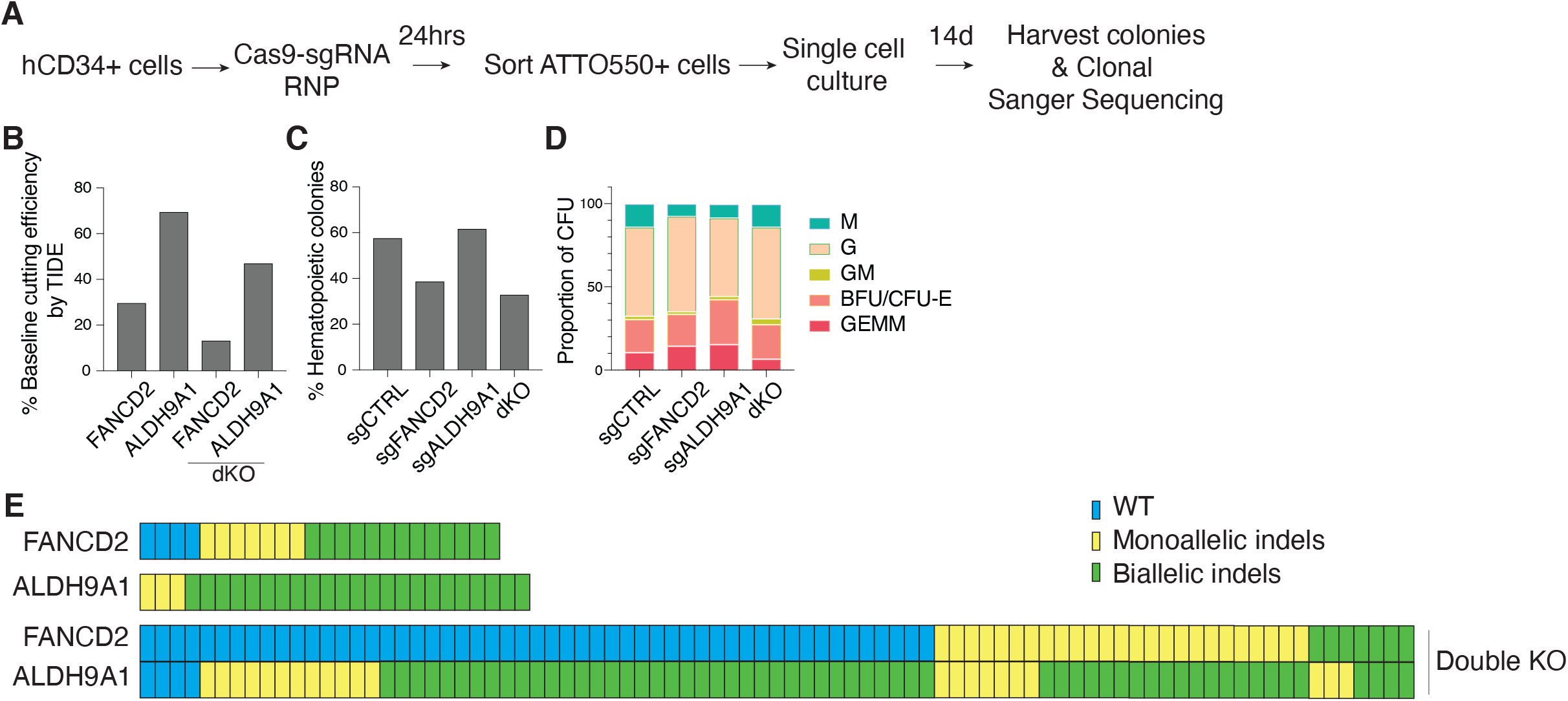
Deficiency of both ALDH9A1 and the Fanconi anemia pathway causes loss of HSPC. A. Schematic of human CD34+ cell editing experiments. Human umbilical cord blood CD34+ cells were edited by electroporation of Cas9-sgRNA-ATTO550 RNP complex. Twenty-four hours later, ATTO550+ cells were single cell sorted onto methylcellulose. Fourteen days later, colonies were scored and harvested for sequencing. B. Baseline editing efficiency was obtained by Sanger sequencing followed by TIDE analysis. C. Percentage of hematopoietic colonies observed per editing condition. D. Proportion of CFU colony subtypes per editing condition. E. Sanger sequencing of each colony per editing condition. The top row shows *FANCD2* sequences of colonies targeted by sgFANCD2 alone. The middle row shows *ALDH9A1* sequences of colonies targeted by sgALDH9A1 alone. The bottom row shows *FANCD2* (upper) and *ALDH9A1* (lower) sequences of colonies targeted by both sgFANCD2 and sgALDH9A1. The frequency of biallelic double KO colony was lower than expected based on baseline editing efficiency assuming *FANCD2* and *ALDH9A1* editing event is independent from each other (observed to expected ratio 0.33). Loss-of-function (LOF) indel was defined as frameshift indels of any size or in-frame indels ≥ 18bp. WT sequence is shown in blue, monoallelic LOF indels in yellow, and biallelic LOF indels in green. dKO, double knockout (i.e., targeted by both sgFANCD2 and sgALDH9A1); M, CFU (colony forming unit)-monocyte/macrophage; G, CFU-granulocyte; GM, CFU-granulocytes/macrophage; BFU-E, burst forming unit-erythroid; CFU-E, colony forming unit-erythroid; GEMM, CFU-granulocyte, erythrocyte, monocyte, megakaryocyte.

### *Fanca^−/−^Aldh9a1^−/−^* mice develop mild hematopoietic phenotypes but show an increased incidence of ovarian tumors with aging

To examine the *in vivo* hematopoietic phenotypes due to increased endogenous aldehydes we expect to form in the absence of ALDH9A1, we generated *Fanca^−/−^Aldh9a1^−/−^* (dKO) mice. These mice were born at the Mendelian ratio without significant anomalies except for small size compared to their littermates and frequent eye abnormalities, which have been previously occasionally observed in FA mouse models^11,27^ (Figure 4A-B; Table S3). Overall survival of the dKO mice was significantly shorter than the WT mice and showed a trend toward shorter survival when compared with *Fanca^−/−^* or *Aldha9a1^−/−^* mice (Figure 4C).

**Figure 4.**
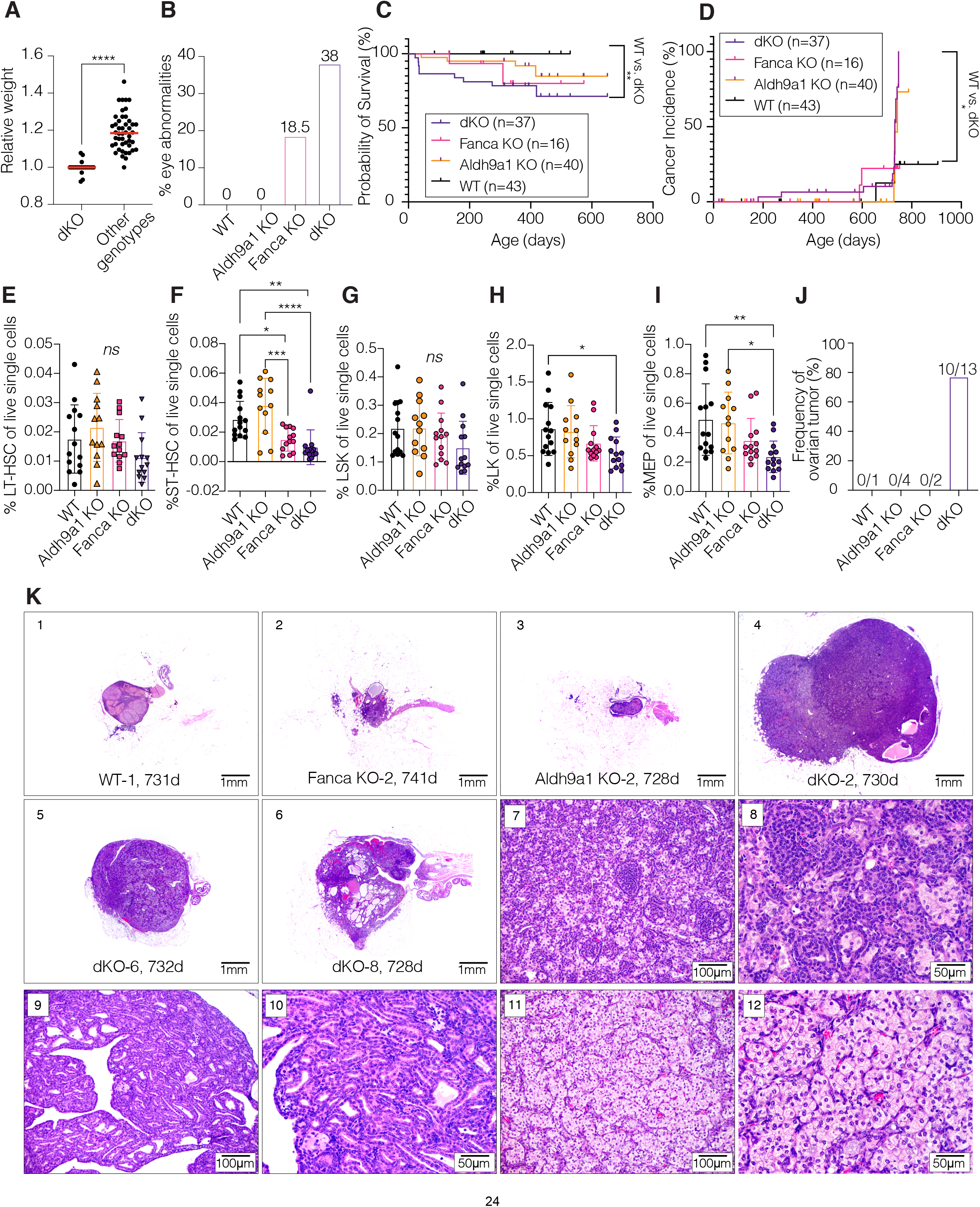
*Fanca^−/−^Aldh9a1^−/−^*mouse model shows mild hematopoietic phenotypes and increased solid tumor incidence with aging. A. Relative weight amongst same-sex littermates. Earliest available weight record was used once in lifetime per mouse between 6 and 10 weeks of age, and normalized to that of same-sex dKO littermates (dKO n=19; other genotypes n=46). Unpaired t test was performed (****, *p* < 0.0001). B. Frequency of eye abnormalities, such as corneal opacity, microphthalmia and anophthalmia (WT n=53, Aldh9a1 KO n=87, Fanca KO n=27, dKO n=50). C. Kaplan-Meier survival curve of four genotype groups. Log-rank test was performed between genotypes (**, *p*=0.0073). D. Kaplan-Meier cancer incidence curve of four genotype groups. Log-rank test was performed between genotypes (*, *p*=0.0171). E-I. Bone marrow aspiration was performed in mice of 8- to 12-weeks of age. Frequency of LT-HSC (E), ST-HSC (F), LSK (G), LK (H) and MEP (I) of live single cells from fresh bone marrow aspirate samples (WT n=14, Aldh9a1 KO n=12, Fanca KO n=13, dKO n=13). One-way ANOVA followed by Tukey’s multiple comparison test was performed (*ns*, not significant; *, *p*<0.05; **, *p*<0.005; ***, *p*<0.0005;****, *p*<0.0001). J. Frequency of ovarian tumor among female mice underwent complete necropsy (age; 9 to 24 months). K. Ovarian histology with Hematoxylin and Eosin stain. 1-3) age-related atrophy of the ovary: 1) WT-1, 2) Fanca KO-2, 3) Aldh9a1 KO-2, 20x magnification. 4) ovarian sex cord stromal tumor, mixed subtype (dKO-2, 20x magnification), a well-demarcated, solid, multilobular, and moderately heterogeneous nodule diffusely replaces the ovarian parenchyma. 5) ovarian sex cord stromal tumor, luteoma subtype (dKO-6, 20x magnification), a well-demarcated, solid, and predominantly homogeneous nodule diffusely replaces the ovarian parenchyma. 6) ovarian tubulostromal adenoma (dKO-8, 20x magnification), a well-demarcated heterogeneous nodule contiguous with the surface epithelium diffusely replaces the ovarian parenchyma and slightly extends into the hilus. Variably cystic tubular structures are interspersed with solid areas of sex cord stromal cells. 7-8) higher magnification of ovarian sex cord stromal tumor, mixed subtype (dKO-2, 200x and 400x magnification respectively), composed of a mixture of sex cord stromal cells at various degrees of differentiation. 9-10) tubulostromal adenoma (dKO-14, 200x and 400x magnification respectively), composed of variably dilated tubules lined by cuboidal epithelium intermixed with small packets of sex cord stromal cells at various degrees of differentiation. 11-12) ovarian sex cord stromal tumor, luteoma subtype (dKO-11, 200x and 400x magnification respectively), composed of packets of highly luteinized polygonal cells supported by a mild fibrovascular stroma. dKO, double knockout (*Fanca^−/−^Aldh9a1^−/−^*); LT-HSC, long-term hematopoietic stem cell; ST-HSC, short-term hematopoietic stem cell; LSK, linage-negative/Sca-1-positive/cKit-high population; LK, lineage-negative/cKit-high population; MEP, megakaryocyte-erythroid progenitor.

dKO mice had the lowest platelet counts compared with WT, *Fanca^−/−^*or *Aldh9a1^−/−^* control mice, but a statistically significant differences was observed only between *Aldh9a1^−/−^* and dKO mice (Figure S2C). Total white blood counts and hemoglobin levels were similar between groups (data not shown). Long-term HSC (LT-HSC), short-term HSC (ST-HSC), lineage-negative, Sca1-positive, cKit-high (LSK), lineage-negative, cKit-high (LK) and megakaryocyte-erythroid population (MEP) were also lowest in dKO mice, but differences were not statistically significant between *Fanca^−/−^* and dKO mice (Figure 4E-I; S2A-B).

However, the effects of ALDH9A1 deficiency in *Fanca^−/−^* mice were revealed with aging. Of the seventeen complete necropsies of aged *Fanca^−/−^Aldh9a1^−/−^* mice (13 females and 4 males; age range: 9-24 months), we observed an increased solid tumor incidence compared to other control groups: 15 of the 17 mice were found to have solid tumors, 10 out of 13 female aged mice had ovarian tumors, and an additional mouse had bilateral ovarian sex cord stromal hyperplasia, which was considered most likely a preneoplastic state (Figure 4D, J-K; Table S4). Ovarian tumors were not observed in any other female mice from control genotypes (Figure 4J).

### The loss of ATP13A3 rescues synthetic lethality of *FANCD2^−/−^ALDH9A1^−/−^* cells

Previous work has implicated ALDH9A1 in the detoxification of aminoaldehydes *in vitro*^28^. However, the *in vivo* substrates of the enzyme have not been identified. To experimentally determine the pathways that created metabolites detoxified by the ALDH9A1, we performed a suppressor screen using the *FANCD2^−/−^ ALDH9A1^−/−^* cells (Figure 5A). To derive *FANCD2^−/−^ALDH9A1^−/−^*clones, we performed nucleofection of Cas9-sgALDH9A1-RNP, and 432 clones were screened from each of *FANCD2^−/−^* c#1 and c#2. Most of clones did not survive and biallelic ALDH9A1 KO was confirmed only in 18 and 5 clones from *FANCD2^−/−^* c#1 and c#2, respectively (Figure S3A-B). A third of those biallelic KO clones failed to expand. Compared with 88-89% indels of ALDH9A1 after nucleofection, the frequency of surviving biallelic dKO clones was extremely low (2.5% and 0.9% for *FANCD2^−/−^* c#1 and c#2, respectively) confirming that *ALDH9A1* knockout is deleterious when combined with FANCD2 deficiency. The clones that survived, most likely retained some ALDH9A1 function. These presumed hypomorphic dKO clones were used to perform a CRISPR screen with the metabolism library, this time looking for guides that would lead to better growth of *FANCD2^−/−^ALDH9A1^−/−^* cells (Figure 5A-B, S3C-D).

**Figure 5.**
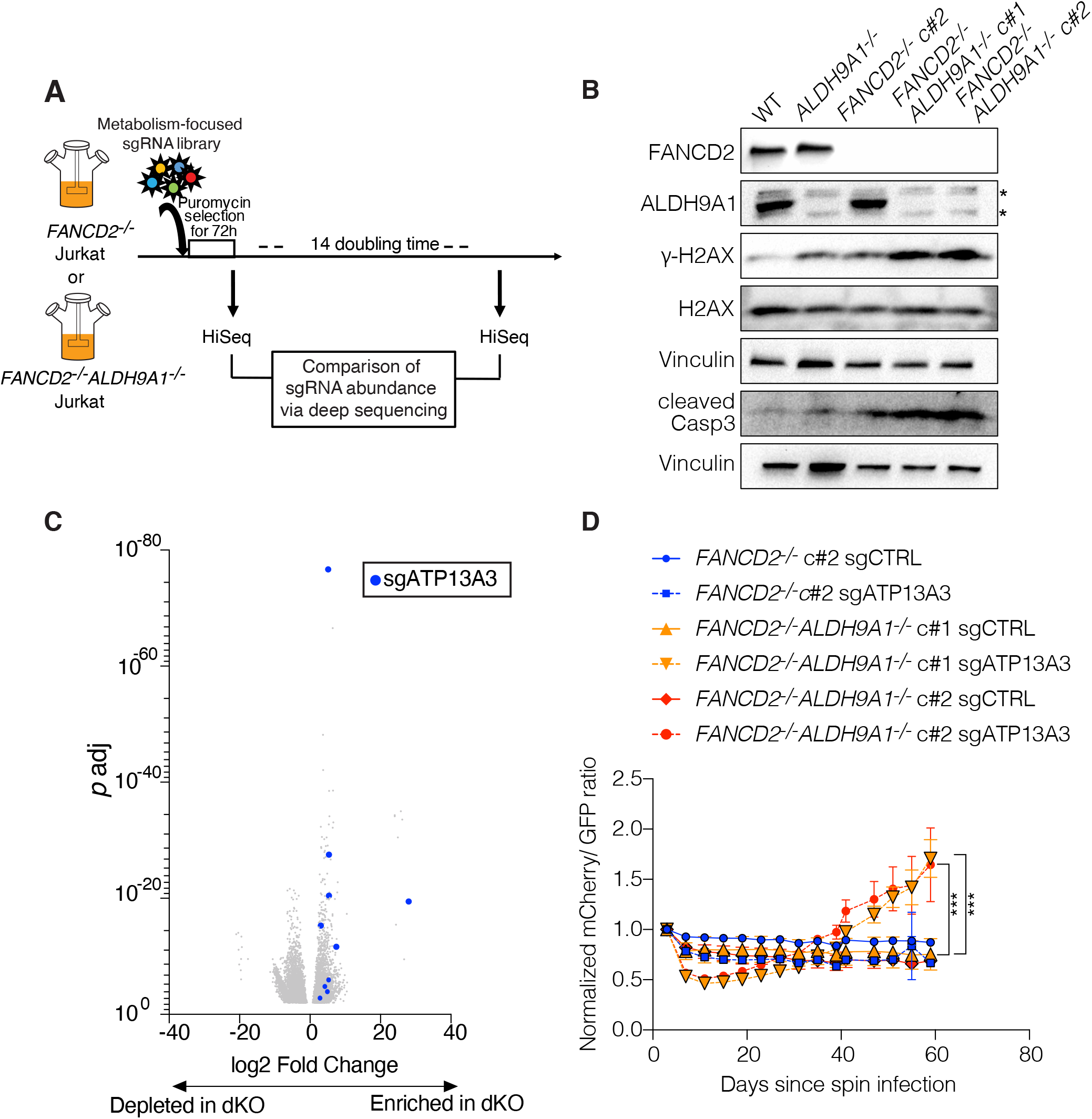
Deficiency of ATP13A3, a polyamine transporter, rescues synthetic lethality of *FANCD2^−/−^ ALDH9A1^−/−^* cells. A. Schematic summary of the suppressor metabolism-focused CRISPR knockout enrichment screen. *FANCD2^−/−^* and *FANCD2^−/−^ALDH9A1^−/−^*Jurkat cells were infected with lentivirus containing metabolism-focused sgRNA library in biological triplicates at MOI of 0.3. sgRNA representation of 500x was maintained throughout the experiment. sgRNAs enriched preferentially in *FANCD2^−/−^ALDH9A1^−/−^* Jurkat cells after 14 population doubling were examined to determine genetic perturbations that rescue synthetic lethality. B. FANCD2, ALDH9A1, gamma-H2AX, H2AX and cleaved Caspase 3 Western blot of Jurkat clones used in the experiments. Vinculin was used as a loading control for each membrane. C. A volcano plot was generated with log2 fold change (log2FC) value and *p* adj value between *FANCD2^−/−^* and *FANCD2^−/−^ALDH9A1^−/−^*at the sgRNA level. Only sgRNAs with *p* <0.01 were used to generate this volcano plot. Ten out of ten sgATP13A3s were significantly enriched in *FANCD2^−/−^ALDH9A1^−/−^* condition and marked as blue dots. D. A competition assay using *FANCD2^−/−^* and two independent *FANCD2^−/−^ALDH9A1^−/−^* clones. The ratio of mCherry/GFP normalized to baseline was progressively increasing in *FANCD2^−/−^ALDH9A1^−/−^*clones with mCherry-sgATP13A3, compared with *FANCD2^−/−^ALDH9A1^−/−^*clones with mCherry-sgCTRL (non-targeting). There were no differences between mCherry-sgALDH9A1 and mCherry-sgCTRL in FANCD2^−/−^ cells. One-way ANOVA followed by Tukey’s multiple comparison test was performed for the last time point. (***, *p*=0.0003) Two independent experiments were performed.

Results of that screen are shown in table S5-6 and Figure 5C. Only loss of one gene, *ATP13A3*, improved the growth of *FANCD2^−/−^ALDH9A1^−/−^*cells when perturbed with multiple guide RNAs (Figure 5C, S3E). sgRNAs targeting *ALDH9A1* were our second top hit enriched in *FANCD2^−/−^ALDH9A1^−/−^*cells. This suggests that targeting of *ALDH9A1* mutant cells might have restored the reading frame and generated partially functional proteins that promoted the survival of *FANCD2^−/−^ALDH9A1^−/−^* cells (Figure S3E).

To validate the rescue of *FANCD2^−/−^ALDH9A1^−/−^* cells by *ATP13A3* knockout, we performed an *in vitro* fluorescence-based competition assay as described earlier using *FANCD2^−/−^* and two independent *FANCD2^−/−^ALDH9A1^−/−^* clones. As expected, we observed expansion of cells with mCherry-sgATP13A3 compared to GFP-sgCTRL only in the two *FANCD2^−/−^ALDH9A1^−/−^*clones, but not in *FANCD2^−/−^* clone or cells with mCherry-sgCTRL (Figure 5D).

### *ALDH9A1* missense variants identified in FA patients are associated with decreased protein expression

To determine whether ALDH9A1 deficiency or any other ALDH or ADH germline variants could modify the clinical presentation of FA patients, we performed targeted DNA sequencing for all known ALDH and ADH superfamily genes in a cohort of subjects with early (n=165) or late (n=60) onset of disease (Figure 6A). In the 225 subjects whose DNA was sequenced, we identified 62 germline variants that were present in the early-onset cohort, but not in the late-onset cohort (Table S7). Among these variants, there were 6 indels, 26 missense variants with a CADD score > 14 (observed in 25 subjects), and 13 missense variants with a CADD score > 20 (observed in 16 subjects). The most recurrent variant among those with a CADD score > 14 was the *ALDH2*2* variant (rs617; CADD score=35), a known disease modifier^15^, observed in 5 subjects (4 heterozygous and 1 homozygous) of East Asian descent. We identified 7 *ALDH9A1* variants in 9 subjects across the early- and late-onset cohorts (4%). Six variants were missense (including one previously reported SNP) and one was a splice site mutation (Figure 6B). The previously reported SNP (rs1065756; gnomAD_exome allelic frequency 0.02) was observed in three subjects and is likely benign (not shown in Figure 6A). Four subjects from the early-onset cohort carried *ALDH9A1* missense variants. Subject 1, whose *ALDH9A1* variant was deemed benign, also carried one variant in *ALDH3A2* (CADD score 5.4) and *ALDH1L1* (rs113725621; CADD score 3.6). Two subjects from the late-onset cohort who carried *ALDH9A1* variants developed malignancies.

**Figure 6.**
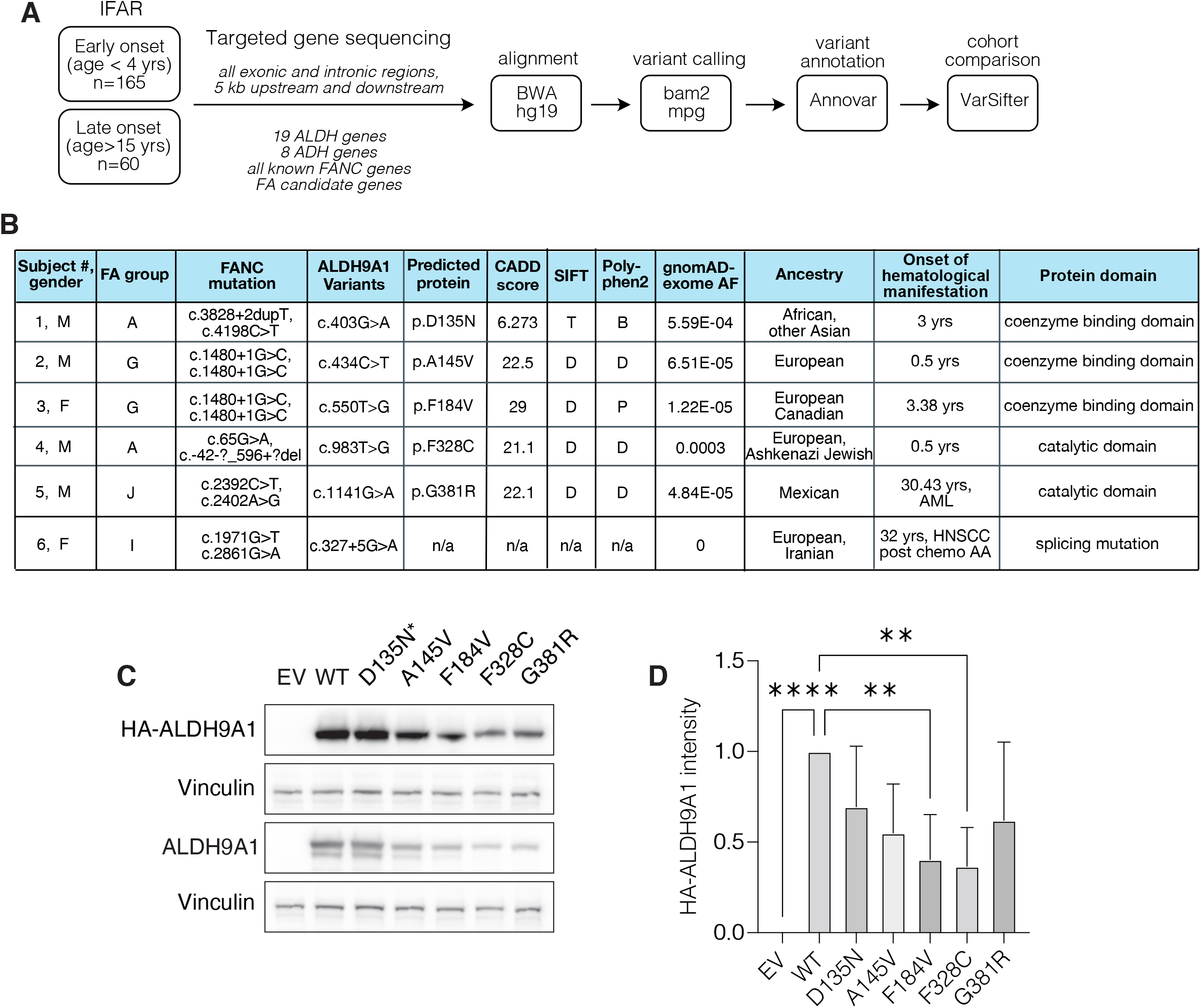
*ALDH9A1* missense variants observed in a subset of Fanconi anemia patients. A. Schematic summary of targeted gene sequencing for all ALDH and ADH genes among International Fanconi Anemia Registry (IFAR) participants with known onset of hematologic manifestations. B. A summary table of six *ALDH9A1* variants observed in patients with Fanconi anemia enrolled in the IFAR (SIFT, T – Tolerated, D – deleterious; Polyphen2, B – benign, D – damaging, P – probably damaging). C. *ALDH9A1* missense variant cDNA was retrovirally over-expressed in *ALDH9A1*^−/−^ Jurkat cells. cDNA expression was confirmed by Western blot using anti-HA antibody and anti-ALDH9A1 antibody. *D135N variant is likely a benign variant given the low CADD score. D. Quantification of baseline HA-ALDH9A1 expression level by ImageJ. HA-ALDH9A1 band intensity was normalized to that of Vinculin loading control. ANOVA followed by Dunnett’s multiple comparison test was performed. (****, *p*<0.0001; **, *p*<0.001) IFAR, International Fanconi Anemia Registry; AF, allelic frequency; M, male; F, female; ND, not determined; n/a, not applicable; HNSCC, head and neck squamous cell carcinoma; AA, aplastic anemia.

To ask if the identified variants expressed lower levels of ALDH9A1, we overexpressed the variants in Jurkat cells. We saw that the missense variants with high CADD scores showed lower expression (Figure 6C-D). Lower protein levels might be partially due to decreased stability as indicated by a trend toward a more rapid decline in protein levels after cycloheximide treatment (Figure S4A-B). More studies of these variants will be necessary to address if they are true modifiers of Fanconi anemia patients.

## Discussion

Endogenous metabolic byproducts, such as reactive aldehydes, are significant sources of DNA damage^29,30^. Inherited deficiency of aldehyde detoxification significantly worsens the outcome of Fanconi anemia in both human and mouse^11–13,15^. Whether ALDH isoenzymes other than ALDH2, and ADH isoenzymes other than ADH5 also provide defense against aldehyde-induced DNA damage had not been investigated. Our study tested 3000 genes in the metabolic pathway in Jurkat cells that were deficient in FANCD2 to identify those that would lead to ICLs when deleted. *ALDH9A1* was the most significantly depleted gene in *FANCD2^−/−^* compared to WT cells. The loss of *ADH5* also caused synthetic lethality in *FANCD2^−/−^* as expected, but to a much lesser degree than the loss of ALDH9A1. The loss of ALDH2 did not cause synthetic lethality in *FANCD2^−/−^* Jurkat cells in pooled screen or single gene KO experiments, which suggests that there exist cell line-specific, and tissue-specific dependencies to the aldehyde detoxification system.

The mouse model of *Fanca^−/−^Aldh9a1^−/−^* (dKO) showed milder phenotypes compared with previously reported *Fancd2^−/−^Aldh2^−/−^* or *Fancd2^−/−^Adh5^−/−^* mouse models^12,13^. While it is possible that the different mouse backgrounds may alter the phenotype severity, we speculate that distinct aldehyde detoxification dependency in mice compared to humans may have contributed to these differences. Analysis of the publicly available RNA-seq dataset confirmed that ALDH2 is the major aldehyde detoxifying enzyme expressed in mouse HSPCs, while human HSPCs express a variety of aldehyde detoxifying enzymes, including ALDH9A1, at much higher levels than ALDH2, raising the possibility that ALDH9A1 plays more important roles in human hematopoiesis than in the mouse (Figure S2D). Increased solid tumor incidence in aged dKO mice compared with control genotypes suggest that the increased basal levels of endogenous DNA damage over their lifetime can contribute to tumorigenesis. This is similar to what is observed in FA patients with hypomorphic *FANC* mutations, which often present with solid tumors later in life with apparently normal bone marrow function until being challenged by chemotherapy^31,32^. These findings suggest that the level and duration of DNA damage, determined by an individual’s DNA repair efficiency as well as aldehyde detoxification capacity, contribute to FA patient’s clinical course – early-onset BMF vs. solid cancer later in life. A remaining question is whether variants of *ALDH9A1* could be associated with ovarian or other cancers even if the FA pathway is intact.

The power of genetics brought by pooled CRISPR screen also identified the presumed culprit aldehyde in the ALDH9A1 deficiency. Investigation of enriched genes identified the loss of ATP13A3 as promoting survival of *FANCD2^−/−^ALDH9A1^−/−^*cells. ATP13A3 transports endocytosed polyamines into the cytosol^33^ where they can be metabolized into 3-aminopropanal, which can spontaneously convert to acrolein^34^. Therefore, the loss of ATP13A3 is predicted to decrease the generation of 3-aminopropanal and/or acrolein by decreasing intracellular polyamines, leading to decreased DNA damage and rescue of *FANCD2^−/−^ ALDH9A1^−/−^* cells. The presence of these aldehydes under different conditions needs to be investigated in future studies.

Approximately 14.5% of the early-onset cohort carried a variant with a CADD score > 20 (n=16) or an indel (n=8) in one of the 19 *ALDH* or 8 *ADH* genes, raising a possibility that broader ALDH and ADH superfamily genes contribute to clinical outcome in FA. Identification of the role of each individual variant in the FA pathogenesis will require further investigation. We also identified a subset of FA patients carrying missense or splice site variants in *ALDH9A1*. Missense variants showed decreased protein levels, which could affect the detoxification capacity of ALDH9A1. Whether this decrease in ALDH9A1 protein expression results in poor patient outcomes will require a larger patient cohort.

In summary, we identified ALDH9A1 deficiency as a previously unrecognized source of endogenous DNA damage. Combined deficiency of ALDH9A1 and FANCD2 caused synthetic lethality due to increased DNA damage and apoptosis. Loss of ATP13A3 rescues *FANCD2^−/−^ALDH9A1^−/−^* cells presumably by decreasing substrates for 3-aminopropanal, which requires ALDH9A1 for its metabolism. Future studies are needed to determine whether enhancement of ALDH9A1 activity or inhibition of ATP13A3 activity could be used to decrease DNA damage load in cells from FA patients and in the general population.

## Methods

### CRISPR knockout screening

A metabolism-focused CRISPR screening was performed as previously described^21^. In short, a pooled sgRNA library with LentiCRISPR v1 vector was used to produce lentivirus by transfection of HEK293T (ATCC) using Trans-IT (Mirus Bio) according to the manufacturer’s protocol. Multiplicity of infection (MOI) was estimated using Jurkat cells by CellTiter-Glo assay (Promega). To ensure integration of only one virus into one cell, lentivirus containing sgRNA library was used at MOI less than 0.3. The volume of virus-containing supernatant to achieve 30% survival after puromycin selection was determined in the virus titration experiments as described above. Jurkat cells were infected by spin infection at 2200 RPM at room temperature for one hour in 6-well plates, containing 2 million cells with polybrene 8ug/mL and lentivirus in a total volume of 2mL per well.

For the dropout screen, to maintain 1000x representation for each sgRNA in the library throughout the experiment, 100 million cells were infected with the virus per replicate and at minimum 40 million cells were passaged each time and harvested for gDNA isolation. For the enrichment screen, a representation of 500x was maintained, therefore 50 million cells were infected with virus and at minimum 20 million cells were passaged each time and harvested for genomic DNA. The experiments were performed in three independent replicates per genotype, however one of three *FANCD2^−/−^* replicates was stopped early due to contamination in the dropout screen, which was excluded from data analysis. For the enrichment screen, three independent replicates were used for each *FANCD2^−/−^* and *FANCD2^−/−^ALDH9A1^−/−^* clones. One of three *FANCD2^−/−^ALDH9A1^−/−^* replicates was an outlier based on the growth curve and data analysis, therefore it was removed from the final data analysis. However, sgATP13A3 was still the most enriched sgRNA in the excluded sample, albeit to a lesser degree.

Genomic DNA was isolated using Blood and Cell Culture DNA Maxi Kit (Qiagen) for the dropout screen, and the Zymo QuickDNA Plus Midiprep Kit (Zymo Research) for the enrichment screen. sgRNA inserts were amplified with NEBNext® High-Fidelity 2X PCR Master Mix for 21 cycles. PCR purification was performed using AMPure XP (Beckman Coulter). Purified PCR products were pooled in equimolar concentration, and sequenced on a NEXTseq 500 (Illumina) with high output 75bp single read setting.

Sequencing data quality was checked using the ShortRead package in R. Sequencing reads were uniquely aligned to the sgRNA library using the Rsubread Bioconductor package’s align function (non default parameters are; unique=TRUE, nBestLocations=1, type = “DNA”, TH1 = 1), and reads per sgRNA counted using the Rsubread’s featureCounts function. Normalized sgRNA abundance and differential sgRNA enrichment between conditions were calculated using the DESeq2 Biconductor package. Gene level depletion or enrichment were summarized as both the number of sgRNAs per gene showing enrichment or depletion between conditions with an Benjamini-Hockberg adjusted p-value < 0.05 and gene level scores were quantified using the mean-rank gene-set enrichment test implemented in the limma Bioconductor package’s genesettest function. Gene ontology enrichment was performed using the TopGO Bioconductor package.

### *In vitro* competition assay

For the first competition assay described in Figure 1G, sgALDH9A1 was inserted to pLentiCRISPRv2-mCherry (Addgene# 99154), and sgCTRL was inserted to pLentiCRISPRv2-GFP (Addgene# 82416) and pLentiCRISPRv2-mCherry. These plasmids were used to generate lentivirus by transfection of HEK293T using Trans-IT. WT, FANCD2^−/−^ c#1 and c#2 were infected with these viruses independently by spin infection as described above. Two days after spin infection, cells infected with lentivirus containing pLentiCRISPR-GFP-sgCTRL was mixed with cells infected with lentivirus containing either pLentiCRISPRv2-mCherry-sgALDH9A1 or pLentiCRISPRv2-mCherry-sgCTRL. The ratio of mCherry/GFP were obtained for baseline on the day of mixing, and then every 4 days by LSRII. The ratio of mCherry/GFP on subsequent days was normalized to that of the baseline.

For the second competition assay described in Figure 4D, sgATP13A3 was inserted to pLentiCRISPRv2-mCherry (Addgene# 99154). Lentiviruses were produced with one of pLentiCRISPRv2-mCherry-sgATP13A3, pLentiCRISPRv2-mCherry-sgCTRL or pLentiCRISPRv2-GFP-sgCTRL. *FANCD2^−/−^*c#2, *FANCD2^−/−^ALDH9A1^−/−^* c#1 and c#2 were infected with these viruses independently by spin infection as described above. Three days after spin infection, cells infected with lentivirus containing pLentiCRISPR-GFP-sgCTRL were mixed with cells infected with lentivirus containing either pLentiCRISPRv2-mCherry-sgATP13A3 or pLentiCRISPRv2-mCherry-sgCTRL. The ratio of mCherry/GFP were obtained for baseline on the day of mixing, and then every 4 days by Aurora cytometer (Cytek Biosciences). The ratio of mCherry/GFP on subsequent days was normalized to that of the baseline.

### Immunofluorescence

At designated time points, cells were harvested and spun on a microscopic slide using Shandon Cytospin 2 centrifuge. After washing with phosphate-buffered saline (PBS) once, cells were fixed with 3.7% formaldehyde (Sigma) in PBS for 10 minutes at room temperature. After washing with PBS twice, cells were permeabilized with 0.5% Triton X-100 (Sigma) in PBS for 10 minutes at room temperature. After washing with PBS twice, cells were incubated in 5% FBS in PBS for one hour at room temperature for blocking, followed by incubation with primary antibody at 4C overnight. After washing cells three times, secondary antibody was incubated for one hour at room temperature. Then cells were washed three times, once with PBS and mounted with DAPI Fluoromount-G® (SouthernBiotech) and coverslips. Images were obtained using Zeiss Axio Observer A1 microscope and AxioVision 4.9.1. software. Data were analyzed by CellProfiler 3.1.8.

### Mice

The Rockefeller University IACUC approved these studies. *Fanca* mouse strain was a gift from Dr. Markus Grompe at Oregon Health Sciences University, Portland, OR. *Aldh9a1* mouse strain was a gift from Dr. Arthur L. Beaudet at Baylor College of Medicine, Houston, TX. *Fanca^−/−^Aldh9a1^−/−^* mice were generated by crossing *Fanca^+/-^* and *Aldh9a1^+/-^*, or *Fanca^+/-^* and *Aldh9a1^−/−^* mice. These mice were maintained on a mixed background for the project. Pups were tailed and genotyped by post-natal day 21.

For retro-orbital bleeding, mice were put under anesthesia by isoflurane inhalation. Once mice are anesthetized, a drop of 0.5% proparacaine HCL ophthalmic solution (Henry Schein, Inc.) was administered to both eyes to maximize the local anesthetic effect. The blunt end of a capillary tube was inserted into the retro-orbital vein. Maximum of 15% of the circulating blood volume was collected into an EDTA tube. Once the bleeding is stopped, antibiotics ointment was applied to the wounded area to prevent infection. The mice received 300 microliters of normal saline support via subcutaneous injection before resting. Complete blood count with differential was performed at the Laboratory of Comparative Pathology at the Memorial Sloan Kettering Cancer Center.

For bone marrow aspiration procedures, we followed a previously published protocol^35^. Briefly, mice received 0.5 ml sterile saline subcutaneously to compensate for loss of body fluid during the procedure. Then the mice were put under Isoflurane inhalation (3%-5%) anesthesia. Mice were given ophthalmic lubricating ointment to the eyes to prevent drying during the procedure. A preemptive dose of buprenorphine was given for pain. After shaving, hind legs were cleaned by betadine solution swab stick followed by 70% ethanol swab scrubbing. A sterile 27g needle was inserted between the two condyles of the femur and into the shaft. The syringe plunger was gently pulled back, creating negative pressure, while moving the needle back and forth within the bone marrow cavity. Successful aspiration was confirmed visually by the appearance of blood in the top of the needle in the base of the syringe. Once bone marrow is successfully aspirated from the femur, we removed the needle and syringe from the femur. A dose of Meloxicam was given for pain.

Mice were submitted for complete necropsy if they become sick (e.g. significant weight loss, hunched-back appearance, or visible tumor > 1.5cm) or at 2 years of age. Three mice submitted for a complete necropsy at 5 weeks of age were excluded from Table S4 and their main findings were hydrocephalus or malocclusion, which can be seen in the C57BL/6 strain as common spontaneous background lesions, therefore not attributed to genotype effect (Aldh9a1 KO n=1, dKO n=2).

### Human HSC experiment

Human umbilical cord blood units were obtained from New York Blood Center. Cells were incubated with HetaSep (Stem Cell Technologies) at 5:1 ratio (cells 5 parts and HetaSep 1 part) for 1 hour at 37C to enhance RBC aggregation. WBCs in the top layer were collected and spun down for 5 minutes at 2000 RPM. Cells were resuspended in wash buffer (2%FBS/IMDM) followed by Ficoll density gradient separation to isolate mononuclear cells. Isolated mononuclear cells were incubated with ammonium chloride solution (Stem Cell Technologies) for RBC lysis, followed by washing with wash buffer twice. Then cells were incubated with human CD34 Microbeads (Miltenyi Biotec) and Fc receptor blocker in MACS buffer (1% BSA/2mM EDTA/PBS) for 30 minutes on ice. Cells were washed and resuspended in MACS buffer, filtered through 40 micron cell strainer, and then CD34+ cells were selected by LS column. After washing, cells were placed in HSC base medium (IMDM containing 20% BIT (Stem Cell Technologies), 100ng/mL recombinant human (rh) SCF, 100ng/mL rhTPO, 10ng/mL rhFLT3-L, and 20ng/mL rhIL-6; cytokines were purchased from PeproTech) in a hypoxic incubator. Next day, HSCs were counted and used 50,000 to 100,000 cells per nucleofection reaction. Ribonucleoprotein (RNP) complex was made by mixing 1.2uL RNA duplex (annealed crRNA and ATTO550-tracrRNA; purchased from IDT), 1.7uL Alt-R® S.p. Cas9 Nuclease V3 and 2.1uL PBS per reaction. Cells were washed with PBS, and resuspended in P3 4D nucleofector solution (Lonza) including RNP complex and electroporation enhancer (IDT), and transferred to nucleocuvette. After nucleofection using 4D Nucleofector (Lonza), cells were incubated in the hypoxic incubator for 24 hours. After 24 hours, single cell sorting was performed using BD ARIA and a single cell was placed into each well of 96-well plate containing Methocult^TM^ H4034 Optimum. Ten to 14 days later, colony subtype was scored and cells were harvested for gDNA extraction and Sanger sequencing.

### Targeted sequencing experiments

The study participants were enrolled in the International Fanconi Anemia Registry (IFAR; Principal Investigator-A.S.) at the Rockefeller University, following written informed consent and/or assent. The Institutional Review Board (IRB) of the Rockefeller University approved these studies. The Office of Human Subjects Research at the National Institutes of Health and IRB of the National Human Genome Research Institute (NHGRI) approved the study using de-identified DNA samples from The Rockefeller University. We selected those with known hematologic onset (defined by the age onset of at least one of the following: ANC < 1000/uL, Hb < 10g/dL, platelets < 100,000/uL, or development of MDS or AML) and available DNA samples. As an early phase of the sequencing effort, we identified 165 participants who had hematologic onset before 4 years of age (early-onset) and 60 patients who had hematologic onset after 15 years of age (late-onset).

A targeted sequencing approach was used by implementing a custom capture kit (Roche, Inc.) for next-generation sequencing designed for 180 genes in total, including all known FA genes and FA candidate genes, eight alcohol dehydrogenase genes, and 19 aldehyde dehydrogenase genes. The panel design included all intronic regions and 5 kb upstream and downstream of each gene. The sequence data was aligned with human genome build hg19 using BWA^36^, variants were called using the Most Probable Genotype algorithm, bam2mpg^37^. Variant annotation was performed using Annovar^38^. The cohort comparison, identifying variants present in the early onset cohort but absent from the late-onset cohort, was performed using VarSifter^39^.

### *ALDH9A1* missense variant overexpression

Five missense variants were identified from the targeted gene sequencing of DNA samples of IFAR participants. WT ALDH9A1 cDNA plasmid was purchased from OriGene (cat#: 216921). A nonsynonymous missense variant (c.26c>G; p.A9G) was detected which is present in the non-conserved region of the WT cDNA. This variant was deemed a SNP without functional consequences but subsequently corrected and other missense variants were generated by QuickChange Multi Site-Directed Mutagenesis Kit (Agilent). WT and missense variant cDNAs were transferred from pDONR223 to MSCV_IP_N terminal-HA-FLAG vector using Gateway recombination (Thermo Fisher Scientific). These vectors were used to produce retrovirus by transfection of HEK293T (ATCC) using Trans-IT (Mirus Bio) according to the manufacturer’s protocol. ALDH9A1-null Jurkat cells were infected by retrovirus containing either empty vector, WT or missense variants by spin infection as described above and selected by puromycin. HA-ALDH9A1 expression was confirmed by western blot. Cells were treated with cycloheximide (Sigma) for 3 and 6 hours and harvested to assess HA-ALDH9A1 expression levels compared to baseline.

## Acknowledgments

We thank Dr. Markus Grompe at Oregon Health Sciences University and Dr. Arthur L. Beaudet at Baylor College of Medicine for sharing *Fanca* and *Aldh9a1* mouse strains, respectively. We also thank Dr. Svetlana Mazel and other staff at the Flow Cytometry Resource Center at the Rockefeller University, staff at the Laboratory of Comparative Pathology at Memorial Sloan Kettering Cancer Center, and Dr. Xiaoling Zhang at the Ross Flow Cytometry Core at Johns Hopkins University for their assistance. This work was supported by grants from the National Heart Lung and Blood Institute (R01 HL120922 - A.S.; K99 HL150628 – M.J.), National Cancer Institute (R01 CA204127 - A.S.), National Center for Advancing Translational Sciences (UL1 TR001866 - M.J., A.S.), American Society of Hematology Scholar Award (M.J.). A.S. is a HHMI Faculty Scholar. R.R.W. was supported by a NRSA F32 from the NIDDK DK115144. This work is partially supported by NIH S10OD026859 (Ross Flow Cytometry Core). F.X.D., R.R-B., and S.C.C. acknowledge the support from the Intramural Research Program of the National Human Genome Research Institute, NIH.

## Contribution

M.J., I.I., Y.P., T.W., D.K., J.A.D, H.B.C, A.G., M.S. J.K., R.W., S.S., R.N., F.P.L. performed experiments. I.M. performed mouse necropsy and histopathology experiments. M.J., I.I., Y.P. A.S. analyzed the results. T.S.C performed genomic data analysis for the CRISPR pooled screen. A.D.A. established the IFAR. F.X.D., R.R-B., and S.C.C. performed targeted sequencing experiments and analyzed the sequencing data. M.J. and A.S. designed the research, made figures, and wrote the paper. All authors edited and approved the manuscript.

## Disclosure

None

## Supplemental Data for

### Supplemental Methods

#### Cell culture

Jurkat clone E6-1 was purchased from ATCC. Jurkat cells were cultured in Roswell Park Memorial Institute 1640 medium (RPMI 1640, Gibco) supplemented with 10% fetal bovine serum (FBS) (Atlanta Biologicals, GA, USA), 1% Pen Strep (Gibco), 1% GlutaMAX™-I 100X (Gibco).

#### Derivation of *FANCD2−/−* Jurkat clones

sgFANCD2 was cloned into pLentiCRISPRv2 (Addgene: #52961) as previously described^35^. pLentiCRISPR v2-sgFANCD2 was transiently expressed in Jurkat by 4D Nucleofector (Lonza) according to the manufacturer’s protocol. One day after nucleofection, puromycin was started at 2ug/mL and continued for 2 days, after which surviving cells were single-cell cloned. Biallelic targeting of *FANCD2* sequence was confirmed by Sanger sequencing and the FANCD2 protein expression was confirmed to be absent by Western blot.

#### MMC sensitivity assay

WT and *FANCD2−/−* Jurkat cells were treated with increasing doses of MMC after which cells were counted using Z2 Coulter® Particle and Size Analyzer (Beckman Coulter). Number of surviving cells were normalized to that of untreated.

#### Western blotting

Cell lysates were obtained by sonication of cells in 2X Laemmli buffer (Bio-Rad) supplemented with 5% 2-Mercaptoethanol (Sigma Aldrich), boiled at 100C for 10 minutes, then used for Western blot. Cell lysates were separated using either a NuPAGE^TM^ 3-8% Tris-Acetate gel or a 4-12% Bis-Tris gel. After protein gel electrophoresis, proteins were transferred to a methanol-activated Immobilon-P PVDF Membrane (EMD Millipore). After transfer, the membrane was blocked with 5% skim milk in tris-buffered saline with 0.05% Tween 20 (TBST) for 1 hour, followed by incubation with primary antibody either overnight at 4C or 2 hours at room temperature. After washing the membrane with TBST three times, it was incubated with secondary antibody for 1 hour at room temperature. After washing with TBST three times, the membrane was developed using Western Lightening Plus-ECL (Perkin Elmer) and images were obtained by Azure c300 Imaging System (Azure Biosystems).

#### Sanger Sequencing

DNA sequence flanking an expected cut site was amplified using Phusion High-Fidelity PCR Master Mix with GC Buffer (NEB). PCR purification was performed with EXOSAP-IT (Applied Biosystems^TM^). Sanger sequencing of purified PCR product was performed by GENEWIZ (South Plainfield, NJ). Data were analyzed using the ICE analysis tool by SYNTHEGO (https://ice.synthego.com/#/).

#### Flow cytometry

Mouse HSC flow cytometry was performed on freshly isolated bone marrow (Table S8). After incubating with ACK lysis buffer for 10 minutes at room temperature for RBC lysis, cells were washed with PBS and incubated with LIVE/DEAD^TM^ Fixable Near-IR Dead Cell Stain Kit (Invitrogen) for 15 minutes at room temperature. Cells were then washed with FACS buffer (2% FBS/2mM EDTA in PBS), and incubated with mouse HSC antibody cocktail (Table S7) for 30 minutes on ice. After washing with FACS buffer, cells were resuspended in 300uL FACS buffer for acquisition. Flow cytometry was performed using BD LSR- Fortessa. Data were analyzed by using FlowJo v.10.

Annexin V staining was performed using PE Annexin V Apoptosis Detection Kit I (BD Biosciences) according to manufacturer’s instruction and events were acquired by BD LSR-Fortessa.

### Supplemental Tables

**Table S1.** Results of the CRISPR screen in WT and *FANCD2−/−* cells. Gene level results are shown. – see excel spreadsheet

**Table S2.** Results of the CRISPR screen in WT and *FANCD2−/−* cells. sgRNA level results are shown. – see excel spreadsheet

**Table S3.** Frequency of eye abnormalities in *Fanca−/−Aldh9a1−/−* mouse model

| Genotype | Total number examined | Number of both eyes affected | Number of one eye affected | Percentage of any eye abnormalities |
| --- | --- | --- | --- | --- |
| *Fanca*<sup>+/+</sup>*Aldh9a1*<sup>+/+</sup> | 53 | 0 | 0 | 0 |
| *Fanca*<sup>+/+</sup>*Aldh9a1*<sup>-/-</sup> | 87 | 0 | 0 | 0 |
| *Fanca*<sup>-/-</sup>*Aldh9a1*<sup>+/+</sup> | 27 | 0 | 5 | 18.5 |
| *Fanca*<sup>-/-</sup>*Aldh9a1*<sup>-/-</sup> | 53 | 3 | 16 | 38 |

Eye abnormalities included corneal opacity, microphthalmia and anophthalmia.

**Table S4.** Results of mouse necropsies and histologic examination.

| Genotype | Mouse ID | Gender | Age (days) | Solid tumors |
| --- | --- | --- | --- | --- |
| Fanca<sup>-/-</sup><br>Aldh9a1<sup>-/-</sup> | dKO-1 | F | 723 | Hepatocellular carcinoma, lymphoma, bilateral ovarian sex cord stromal tumors |
|  | dKO-2 | F | 730 | Bilateral ovarian sex cord stromal tumors, thyroid follicular adenoma, histiocytic sarcoma |
|  | dKO-3 | F | 730 | None (sex cord stromal hyperplasia in bilateral ovaries) |
|  | dKO-4 | F | 274 | Right ovarian sex cord stromal tumor |
|  | dKO-5 | F | 741 | Lymphoma |
|  | dKO-6 | F | 732 | Left ovarian tubulostromal adenoma; right ovarian sex cord and interstitial cell hyperplasia |
|  | dKO-7 | F | 732 | Bilateral ovarian sex cord stromal tumors; lymphoma |
|  | dKO-8 | F | 728 | Left ovarian tubulostromal adenoma; Right ovarian sex cord stromal tumor, lymphoma |
|  | dKO-9 | F | 605 | Bilateral ovarian tumors, lymphoma |
|  | dKO-10 | F | 736 | Right ovarian sex cord stromal tumor and cystadenoma |
|  | dKO-11 | F | 736 | Right ovarian sex cord stromal tumor |
|  | dKO-12 | F | 454 | None |
|  | dKO-13 | F | 180 | Lymphoma |
|  | dKO-14 | F | 747 | Bilateral ovarian tumulostromal adenoma, lymphoma |
|  | dKO-15 | M | 731 | Lymphoma |
|  | dKO-16 | M | 732 | Hemangiosarcoma |
|  | dKO-17 | M | 732 | Lymphoma |
|  | dKO-18 | M | 151 | None |
|  | dKO-19 | M | 721 | Cutaneous Hemangioma |
| Fanca<sup>+/+</sup><br>Aldh9a1<sup>-/-</sup> | Aldh9a1 KO-1 | F | 730 | Lymphoma, pulmonary adenoma |
|  | Aldh9a1 KO-2 | F | 728 | Histiocytic sarcoma |
|  | Aldh9a1 KO-3 | F | 732 | None |
|  | Aldh9a1 KO-4 | F | 731 | Pituitary gland adenoma (sex cord stromal hyperplasia in bilateral ovaries) |
|  | Aldh9a1 KO-5 | M | 730 | None |
|  | Aldh9a1 KO-6 | M | 677 | None |
|  | Aldh9a1 KO-7 | M | 742 | Hemangiosarcoma |
| Fanca<sup>-/-</sup><br>Aldh9a1<sup>+/+</sup> | Fanca KO-1 | F | 743 | None |
|  | Fanca KO-2 | F | 741 | None |
|  | Fanca KO-3 | M | 730 | None |
|  | Fanca KO-4 | M | 598 | Lymphoma |
|  | Fanca KO-5 | M | 735 | None |
|  | Fanca KO-6 | M | 589 | Lymphoma |
| Fanca<sup>+/+</sup><br>Aldh9a1<sup>+/+</sup> | WT-1 | F | 731 | Splenic Hemangiosarcoma |
|  | WT-2 | M | 655 | Hepatocellular carcinoma |

**Table S5.** Results of the suppressor CRISPR screen in *FANCD2−/−* and *FANCD2−/−ALDH9A1−/−* cells. Gene level results are shown. – see excel spreadsheet

**Table S6.** Results of the suppressor CRISPR screen in *FANCD2−/−* and *FANCD2−/−ALDH9A1−/−* cells. sgRNA level results are shown. – see excel spreadsheet

**Table S7.**
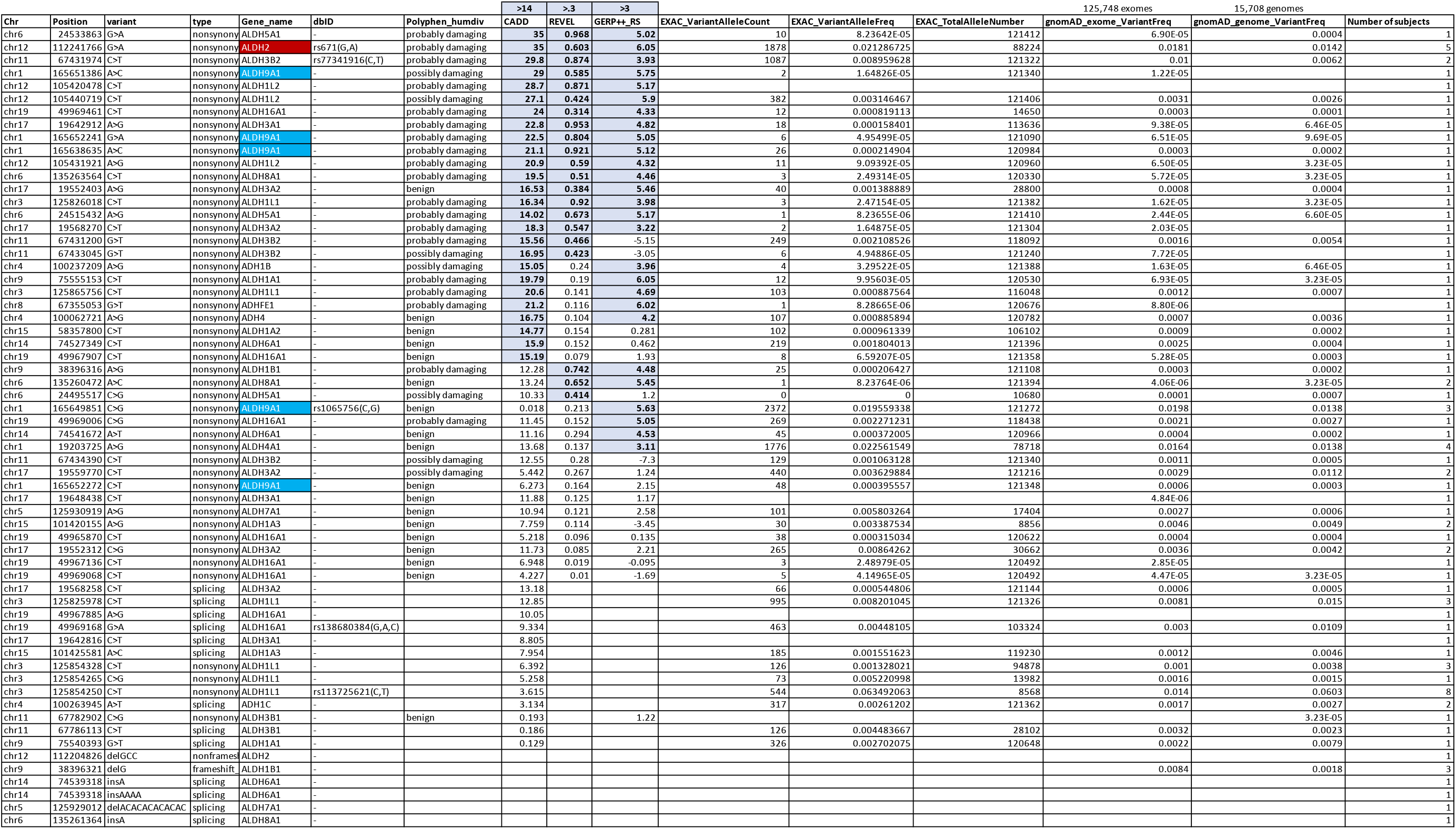
ALDH and ADH variatns identified in subjects with Fanconi anemia. Shown are the variants present in individuals from the early-onset cohort and absent from the late-onset cohort. Coordinates are in accord with human genome build 19

**Table S8.** List of antibodies used in the experiments

**Primary antibodies**
| Antibody | Host | Clone | Vendor | Catalog# |
| --- | --- | --- | --- | --- |
| HA | mouse | 16B12 | BioLegend | 901514 |
| α-tubulin | mouse | DM1A | Sigma | T9026 |
| γH2AX | mouse | JBW301 | Millipore | 05-636 |
| H2AX | rabbit |  | Sigma | PLA0023 |
| 53BP1 | rabbit |  | Novus Biologicals | NB100-304 |
| FANCD2 | rabbit |  | Novus Biologicals | NB100-182 |
| Vinculin | mouse | hVIN-1 | Sigma | V9131 |
| ALDH9A1 | rabbit |  | Abcam | ab224360 |

**Secondary antibodies**
| Antibody | Vendor | Catalog# |
| --- | --- | --- |
| Alexa Fluor 488 goat anti-rabbit IgG (H+L) | Life Technologies | A11008 |
| Alexa Fluor 594 goat anti-rabbit IgG (H+L) | Life Technologies | A11012 |
| Alexa Fluor 488 goat anti-mouse IgG (H+L) | Life Technologies | A11029 |
| Alexa Fluor 488 goat anti-mouse IgG (H+L) | Life Technologies | A11005 |
| Peroxidase AffiniPure Goat Anti-Rabbit IgG (H+L) | Jackson ImmunoResearch Laboratories, INC | 111-035-003 |
| Peroxidase-conjugated AffiniPure Goat Anti-Mouse IgG (H+L) | Jackson ImmunoResearch Laboratories, INC | 115-035-003 |

**Flow cytometry antibodies**
| Marker | Fluorochrome | Clone | Vendor | Cat# |
| --- | --- | --- | --- | --- |
| CD34 | FITC | RAM34 | ThermoFisher | 11-0341-82 |
| c-Kit | APC | 2B8 | ThermoFisher | 17-1171-82 |
| CD16/CD32 | AF700 | 93 | ThermoFisher | 56-0161-82 |
| viability | Near IR |  | ThermoFisher | L34975 |
| Sca-1 | BV421 | D7 | Biolegend | 115904 |
| CD48 | BV510 | HM48-1 | BD Pharmigen | 563536 |
| CD150 | PE | TC15-12F12.1 | Biolegend | 115904 |
| γH2AX | PE-CF594 | N1-431 | BD Pharmigen | 564719 |
| CD3e | BUV395 | 145-2C11 | BD Pharmigen | 563565 |
| CD4 | BUV395 | RM4-5 | BD Pharmigen | 740208 |
| CD8a | BUV395 | 53-6.7 | BD Pharmigen | 563786 |
| B220 | BUV395 | RA3-6B2 | BD Pharmigen | 563793 |
| CD11b | BUV395 | M1/70 | BD Pharmigen | 563553 |
| Gr1 | BUV395 | RB6-8C5 | BD Pharmigen | 563849 |
| Ter119 | BUV395 | Ter-119 | BD Pharmigen | 563827 |
| CD41 | BUV395 | MWReg30 | BD Pharmigen | 564056 |

**Figure S1.**
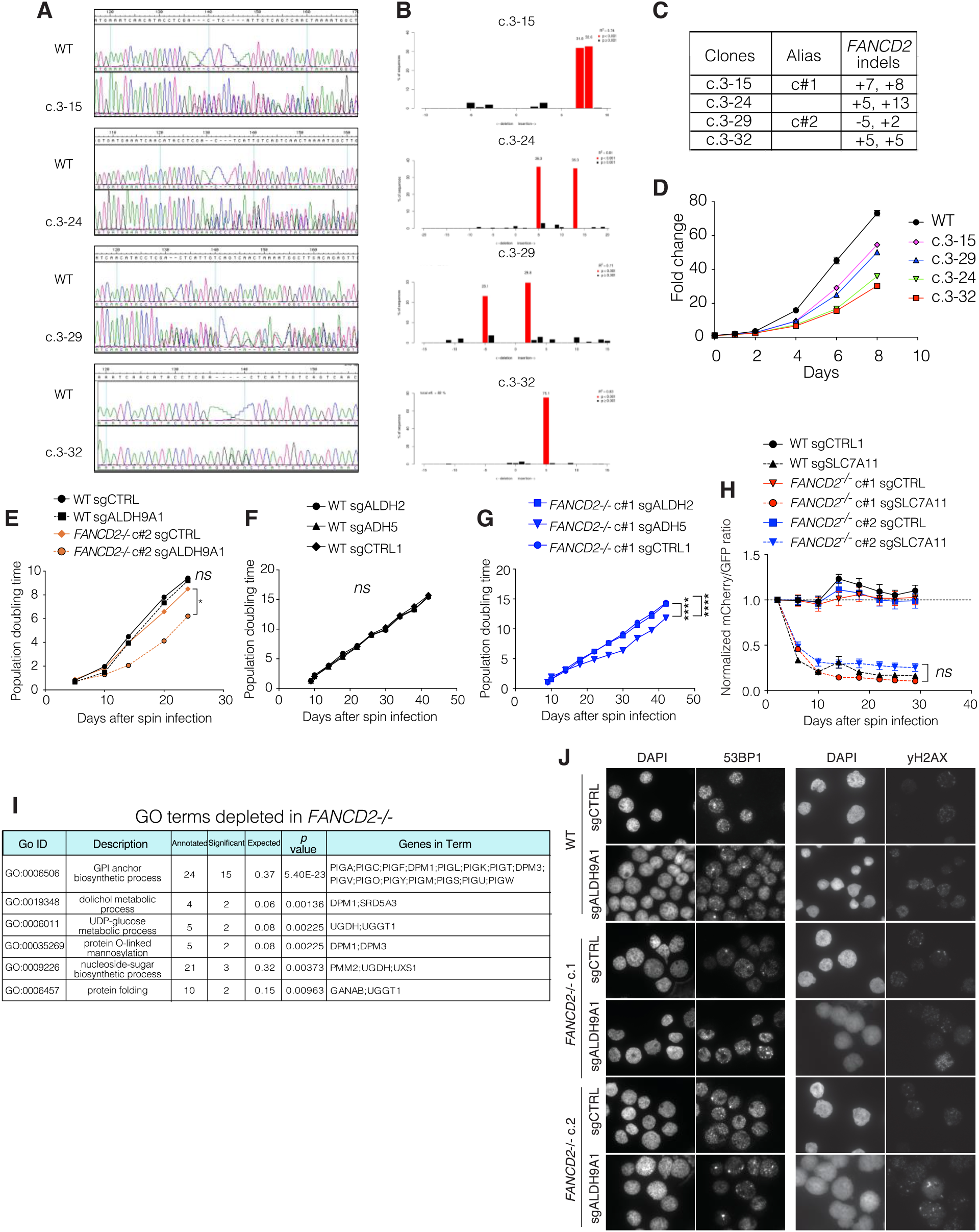
Excessive reactive aldehydes caused by ADH5 or ALDH9A1 deficiency cause synthetic lethality in FANCD2-deficient cells by increased DNA damage. A. Genomic sequence of four *FANCD2−/−* clones compared to that of WT. B. Indel of *FANCD2* gene was inferred by Tracking of Indels by DEcomposition (TIDE) analysis (http://shinyapps.datacurators.nl/tide/). C. A summary table of indels for four *FANCD2−/−* clones. D. A growth curve of WT and four *FANCD2−/−* clones (mean +/- SEM). E-G. WT and *FANCD2−/−* Jurkat cells were infected by lentivirus containing Cas9 and sgRNA. After puromycin selection for 3 days, cumulative population doublings were calculated. WT cells showed no differences in population doubling time regardless of sgRNA used. *FANCD2−/−* cells showed lower population doubling time when sgADH5 or sgALDH9A1 were used, but not when sgALDH2 or sgCTRL (non-targeting) were used. One-way ANOVA followed by Tukey’s multiple comparison test was performed for the last time point. (ns, not significant; *, *p*<0.05; ****, *p*<0.0001). H. A competition assay as in 1G using mCherry-sgSLC7A11. Cells infected with mCherry-sgSLC7A11 were significantly outcompeted by cells infected with GFP-sgCTRL in all cell lines tested. Such differences were not observed when mCherry-sgCTRL was used instead. One-way ANOVA test was performed for the last time point for WT-sgSCL7A11, *FANCD2−/−* c#1-sgSLC7A11 and *FANCD2−/−*c#2-sgSLC7A11 (ns, not significant). I. GO terms depleted in *FANCD2*−/−. Multiple genes in the GPI anchor biosynthetic process were depleted in *FANCD2*−/− cells, suggesting disruption of this pathway is detrimental to the survival of *FANCD2*−/− cells. J. Immunofluorescence assay for 53BP1 and gamma-H2AX was performed on bulk KO samples. Quantification is shown in Figure 2E-F. SEM, standard error or mean; GO, gene ontology; GPI, glycosylphosphatidylinositol.

**Figure S2.**
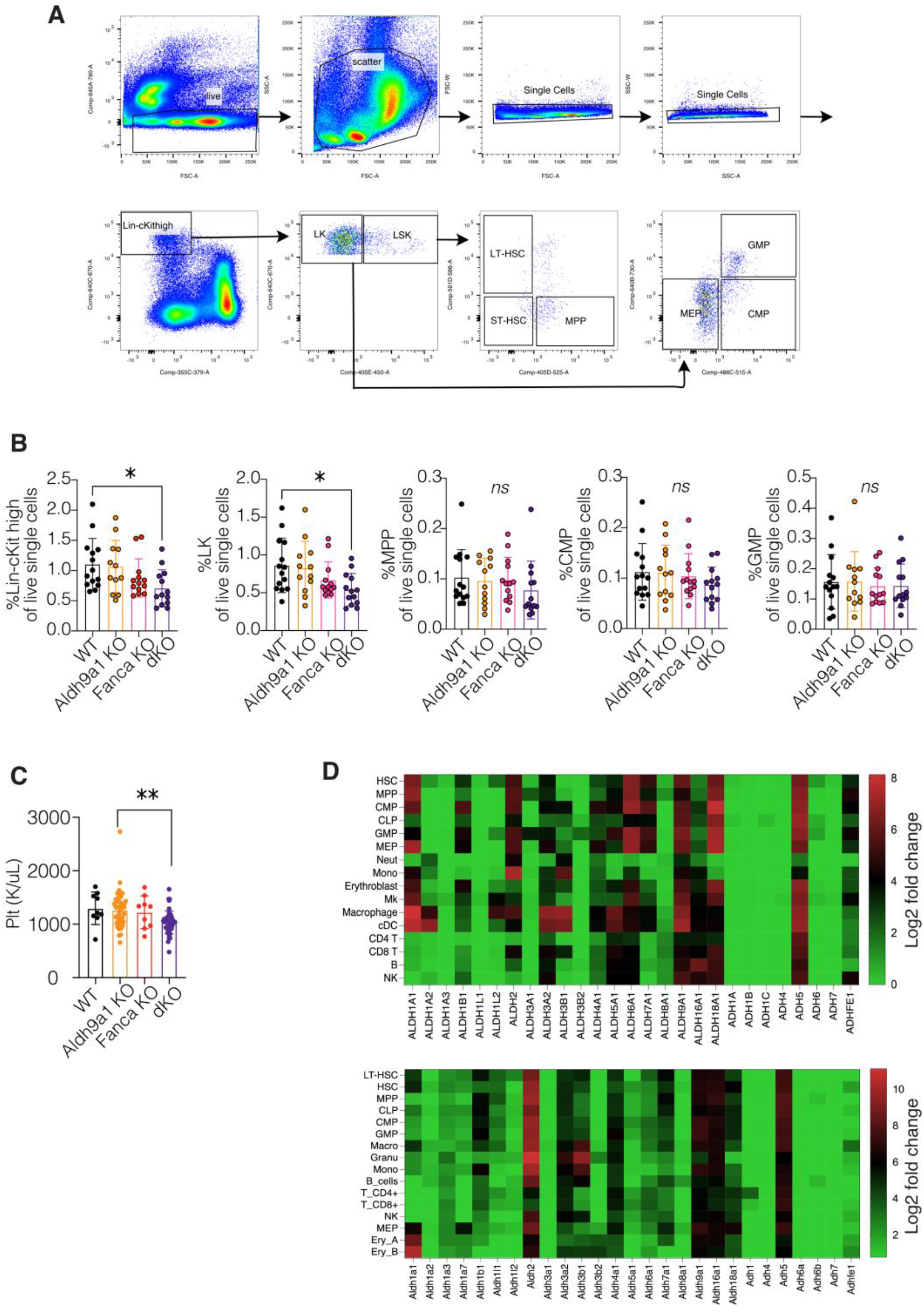
Mouse hematopoiesis shows a predominant ALDH2 expression, compared to human hematopoiesis in which shows higher expression of other ALDH isoenzymes than ALDH2. A. Mouse HSC flow gating scheme is shown. B. Frequency of Lineage-negative/cKit-high population, LK, MPP, CMP and GMP of live single cells from fresh bone marrow aspirate samples (WT n=14, Aldh9a1 KO n=12, Fanca KO n=13, dKO n=13; 8-12 weeks old). One-way ANOVA followed by Tukey’s multiple comparison test was performed (*ns*, not significant; *, *p*<0.05). C. Peripheral blood platelet counts of 9- to 12-month-old mice (WT n=9, Aldh9a1 KO n=44, Fanca KO n=9, dKO n=45). One-way ANOVA followed by Tukey’s multiple comparison test (**, *p*=0.0012). D. Gene expression data for human and mouse ALDH and ADH isoenzymes in hematopoietic cells. Publicly available single cell RNA-seq data were downloaded from BloodSpot.eu and a heatmap was generated using log2 fold change values on Prism GraphPad. LK, Lineage-negative/cKit-high/Sca-1 negative population; MPP, multipotent progenitor; CMP, common myeloid progenitor; GMP, granulocyte-macrophage progenitor.

**Figure S3.**
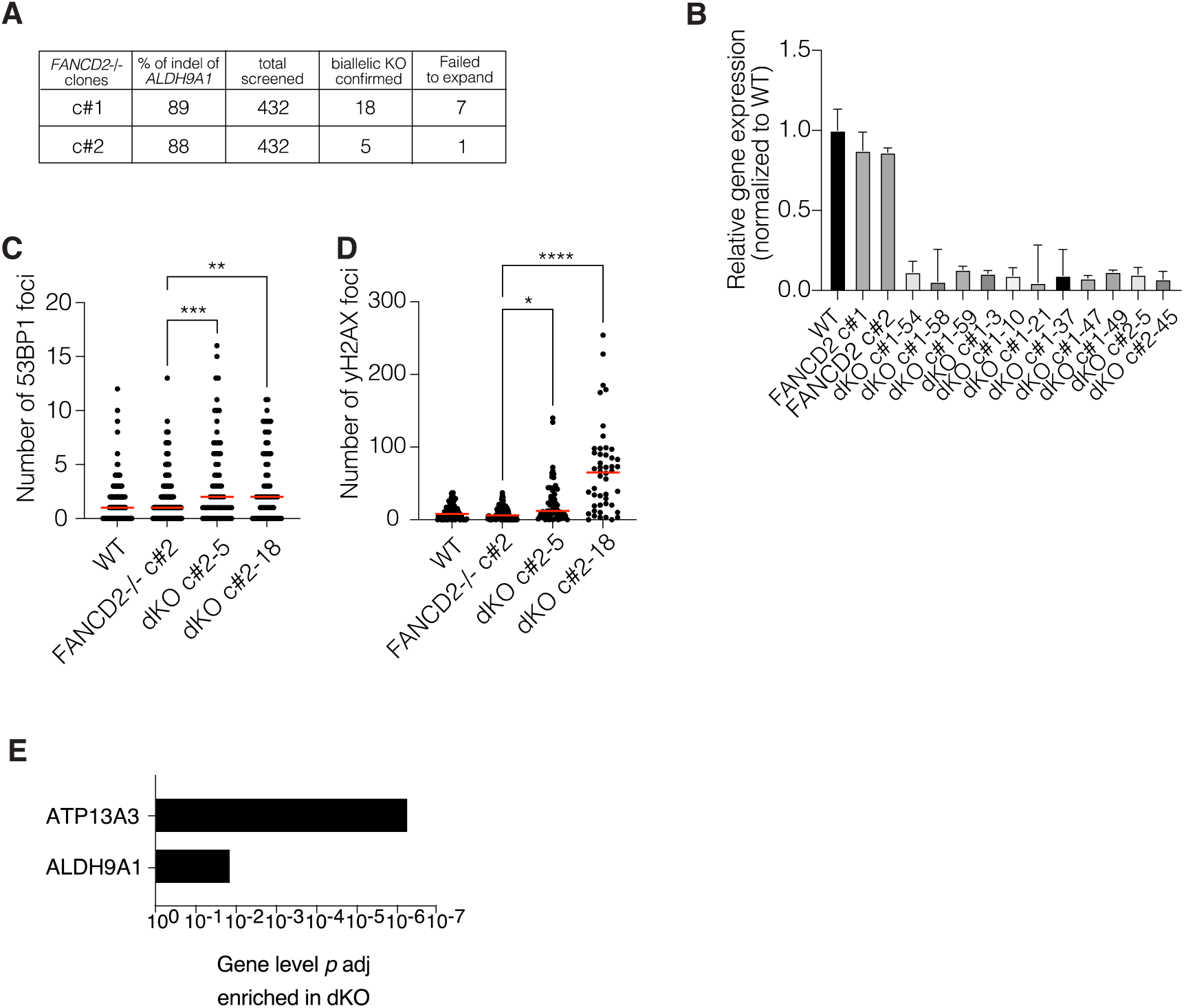
*FANCD2−/−ALDH9A1−/−* clones showed increased DNA damage and poor survival. A. Summary table for single-cell cloning of *FANCD2−/−ALDH9A1−/−* cells. B. ALDH9A1 RT-qPCR results for WT, FANCD2−/− and FANCD2−/−ALDH9A1−/− (dKO) clones. C-D. Numbers of 53BP1 foci per cell (C) and gamma-H2AX foci per cell (D) by immunofluorescence were elevated in two dKO clones compared with their parental *FANCD2−/−* clone. One-way ANOVA followed by Tukey’s multiple comparison test was performed (*, *p*<0.05; **, *p*<0.005; ***, *p*<0.0005;****, *p*<0.0001). E. *p* adj values for genes that were differentially enriched in *FANCD2−/−ALDH9A1−/−* condition (dKO) were calculated by the DESeq2 package and shown as a minus log 10 graph.

**Figure S4.**
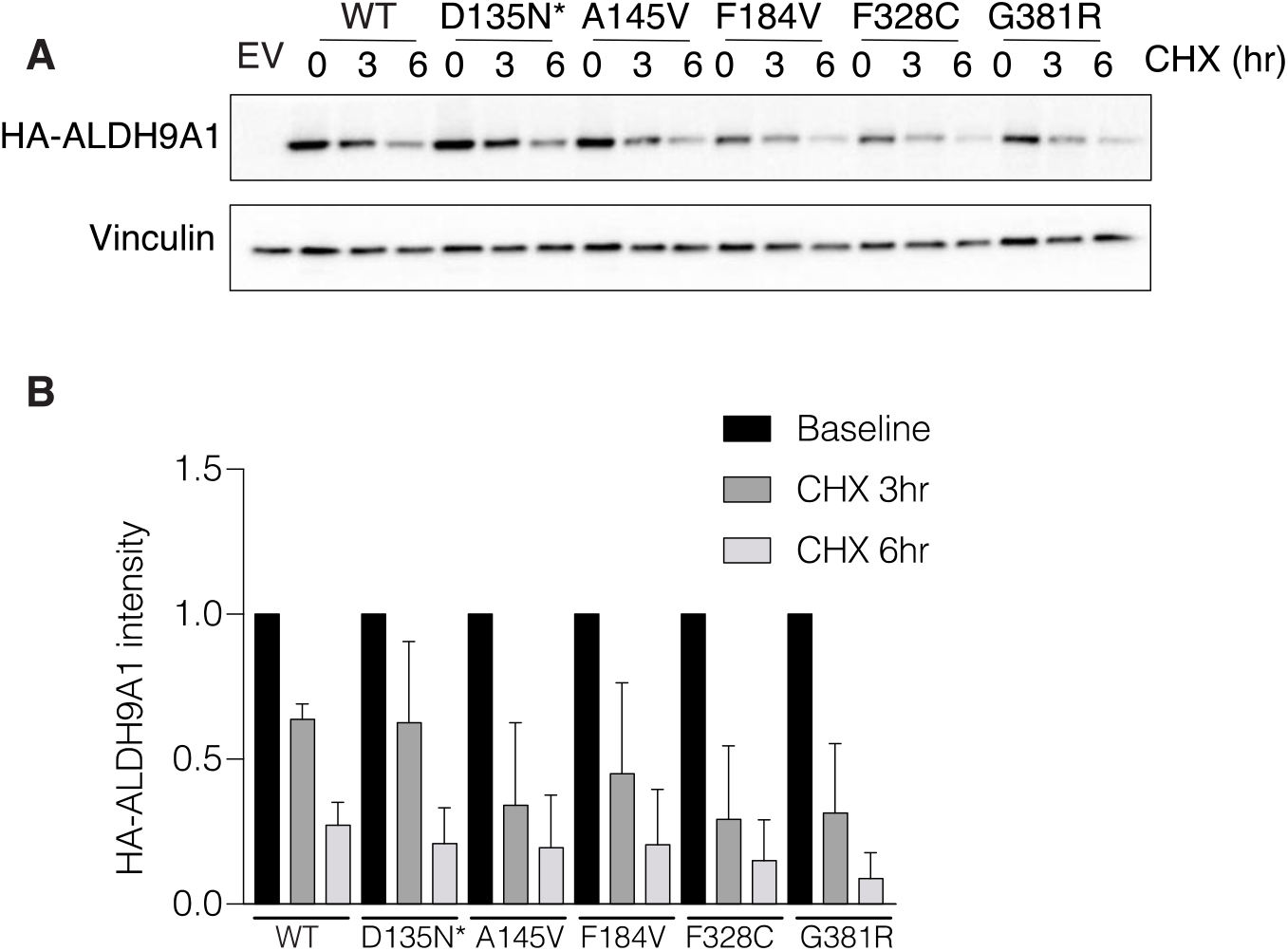
*ALDH9A1* missense variants show decreased protein stability. A. *ALDH9A1*−/− Jurkat cells overexpressing *ALDH9A1* cDNA were treated with cycloheximide (CHX) 50ug/mL for the indicated duration, after which HA-ALDH9A1 expression was assessed by Western blot. B. Quantification of HA-ALDH9A1 expression following CHX treatment (normalized to Vinculin loading control). Differences between missense variants at each time point were not statistically significant and showed a trend only. ANOVA followed by Dunnett’s multiple comparison test was performed.

## Notes

### Competing Interest Statement

The authors have declared no competing interest.

